# EZH2 coordinates inflammatory and pain-associated pathways across joint compartments in osteoarthritis

**DOI:** 10.1101/2025.01.19.633762

**Authors:** Sybille Brochard, Alison Vanlaeys, Mahdia Taieb, Isabelle Richard, Juliette Aury-Landas, Benoit Bernay, Julien Pontin, Celia Seillier, Nina Bon, Baptiste Picart, Claire Pecqueur, Olivier Toutirais, Claire Vinatier, Jérôme Guicheux, Eric Maubert, Véronique Agin, Karim Boumediene, Catherine Bauge

## Abstract

Enhancer of zeste homolog 2 (EZH2), a histone methyltransferase responsible for H3K27 trimethylation, has emerged as a potential therapeutic target in osteoarthritis (OA). However, its contribution to the multicellular mechanisms driving joint degeneration and pain remains poorly understood. Here, we investigated the effects of pharmacological EZH2 inhibition in a pain-relevant murine OA model and explored its cellular and molecular consequences across OA-associated cell populations. OA was induced in mice by intra-articular monosodium iodoacetate (MIA) injection followed by local administration of the EZH2 inhibitor EPZ-6438. Joint pathology and pain-related behaviors were assessed by histological and functional analyses. Mechanistic studies were performed in primary human OA fibroblast-like synoviocytes and bone marrow-derived cells using targeted gene expression analyses, proteomics and ChIP-seq approaches. EZH2 inhibition reduced cartilage damage, synovial inflammation and pain-related behavioral alterations in vivo. In OA synoviocytes, EPZ-6438 decreased the expression of inflammatory, catabolic and pain-associated mediators while promoting autophagy-related responses. Proteomic and ChIP-seq analyses revealed EZH2-dependent regulation of inflammatory pathways, cellular homeostasis and neuronal-associated processes, including axon guidance-related pathways. ChIP-seq further identified inflammation-dependent EZH2 recruitment to promoters of neurodevelopmental regulators, including PAX6, suggesting a potential contribution of EZH2 to neuronal-associated mechanisms in OA. In addition, EZH2 inhibition reduced macrophage inflammatory activation and osteoclast differentiation. Together, these findings identify EZH2 as a candidate epigenetic regulator linking inflammatory, neuroimmune and osteoimmune pathways across the osteoarthritic joint. Targeting EZH2 may represent a therapeutic strategy to simultaneously modulate joint inflammation, remodeling and pain-associated pathways.

## Introduction

Osteoarthritis (OA) is the most prevalent chronic joint disease and a major cause of pain and disability worldwide, affecting more than 500 million individuals globally (1). Although historically considered a degenerative disorder primarily characterized by cartilage loss, OA is now recognized as a complex whole-joint disease involving dynamic interactions between articular cartilage, synovium, immune cells, subchondral bone, and peripheral sensory nerves (2,3). This revised disease paradigm highlights the importance of identifying molecular regulators capable of integrating the multiple cellular processes responsible for both structural deterioration and pain development.

Current therapeutic strategies for OA remain largely symptomatic and focus mainly on pain management. No approved disease-modifying osteoarthritis drug is currently available to prevent or reverse disease progression. In advanced stages, joint replacement surgery remains the only effective intervention to restore function in severely affected patients (2). The urgent need for innovative therapeutic approaches has therefore stimulated research aimed at identifying molecular regulators capable of simultaneously targeting joint destruction and pain mechanisms.

Among emerging regulatory mechanisms, epigenetic modifications have gained increasing attention as important contributors to OA pathogenesis. Epigenetic regulation provides a dynamic mechanism allowing cells to adapt their transcriptional programs in response to aging, inflammation, mechanical stress, and metabolic changes. Alterations in DNA methylation, non-coding RNA expression, and histone modifications have been associated with OA development and progression (4,5). In particular, histone methylation at lysine 27 of histone H3 (H3K27me3), a repressive chromatin mark mainly deposited by the histone methyltransferase Enhancer of Zeste Homolog 2 (EZH2), has emerged as an important regulator of cartilage homeostasis.

EZH2, the catalytic component of the Polycomb Repressive Complex 2 (PRC2), is a key epigenetic regulator controlling gene expression through deposition of the repressive H3K27me3 mark. Several studies, including previous work from our group, have demonstrated that EZH2 is activated during cartilage degeneration and contributes to chondrocyte hypertrophy, inflammatory responses, and extracellular matrix breakdown (6–9). However, these observations have mainly focused on cartilage, leaving unresolved whether EZH2 participates in the regulation of other cellular compartments involved in OA pathogenesis. In particular, the contribution of EZH2 to synovial inflammation, immune cell activation, bone remodeling, and OA-associated pain remains poorly understood. This represents an important knowledge gap because pain, rather than structural changes alone, is the primary determinant of clinical burden and patient disability. Increasing evidence indicates that OA pain results from complex interactions between inflammatory mediators, immune cells, synovial alterations, subchondral bone remodeling, and sensory nerve signaling (10–12). Identifying molecular regulators that simultaneously influence these processes could therefore provide new opportunities for disease-modifying and analgesic therapies.

Recent findings suggest that epigenetic regulators may influence immune responses and tissue communication. In particular, EZH2 has been implicated in macrophage activation, inflammatory polarization, and osteoclast differentiation in several pathological contexts, suggesting that it may represent a broader regulator of inflammatory and remodeling responses (13,14). However, whether EZH2 contributes to the coordinated regulation of multiple cellular compartments during OA progression remains unknown.

In this study, we hypothesized that EZH2 represents a central epigenetic regulator extending beyond cartilage degeneration to control inflammatory, immune, and pain-associated pathways within the OA joint. Using a combination of in vivo pharmacological inhibition, human cellular models, proteomic profiling, and genome-wide EZH2 binding analysis, we investigated how EZH2 influences joint pathology across multiple cellular compartments, including synoviocytes, macrophages, and osteoclasts.

## Results

### EZH2 inhibition protects joint integrity and alleviates pain-related alterations in experimental OA

Previous studies have established EZH2 as a regulator of cartilage degeneration in surgically induced OA models, but its contribution to OA-associated pain remains unclear. To address this question, we used the monosodium iodoacetate (MIA)-induced OA model, which combines progressive joint structural damage with robust pain-related behavioral alterations.

OA-like pathology was induced by intra-articular injection of MIA, and mice received local administration of the selective EZH2 inhibitor EPZ-6438 at days 2, 7, and 30 following OA induction (Figure 1a). MIA injection resulted in progressive cartilage degradation, synovial alterations, mechanical hypersensitivity, and impaired weight bearing. In contrast, EPZ-6438-treated animals displayed reduced structural and functional alterations.

**Figure 1:**
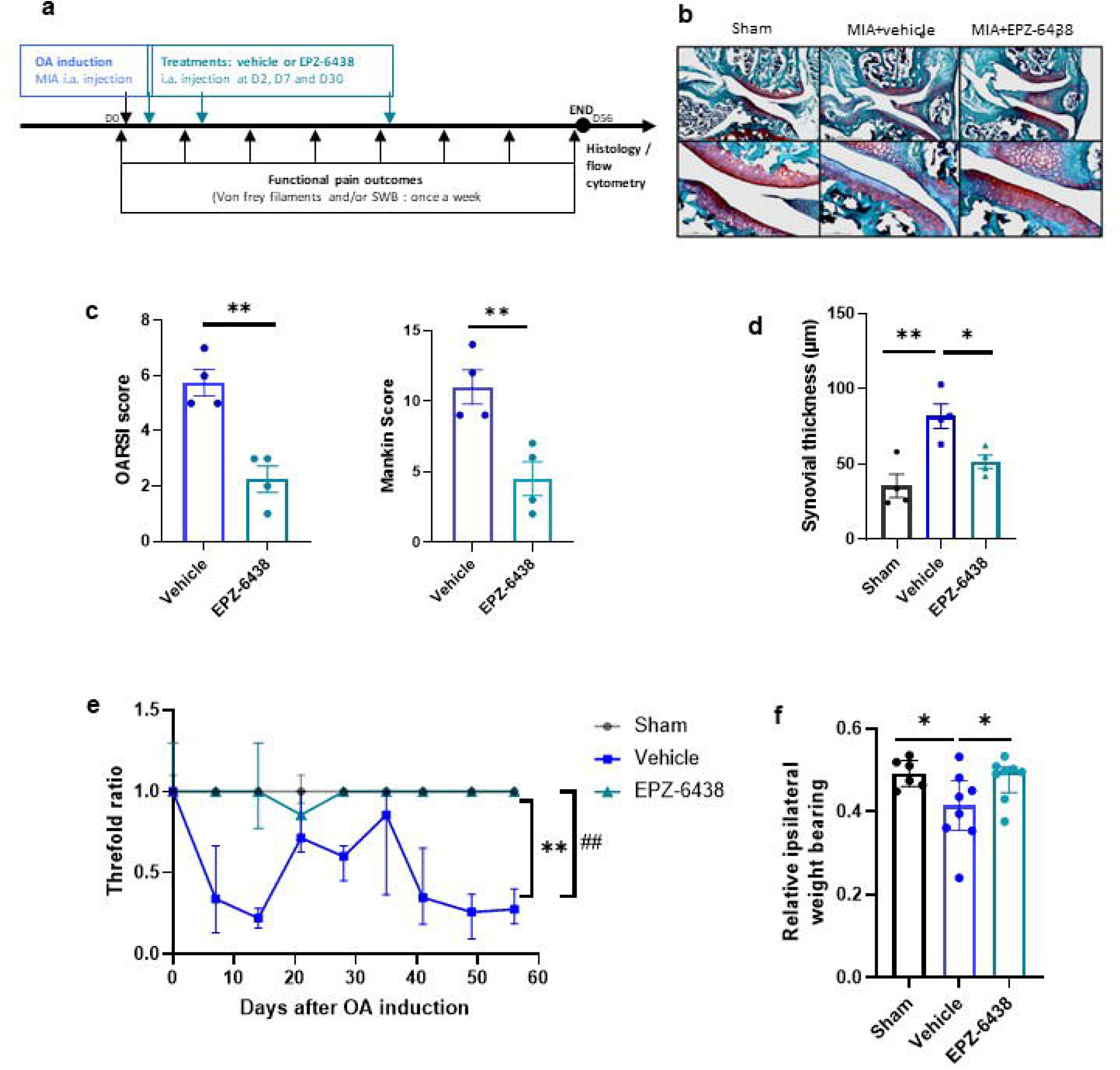
EZH2 inhibition reduces OA progression and pain in a murine model of osteoarthritis. Schematic representation of the experimental design (a). Osteoarthritis was induced by intra-articular injection of monosodium iodoacetate (MIA, 0.75 mg diluted in 10 μL of sterile saline solution). Sham animals received intra-articular injection of vehicle. At days 2, 7 and 30 after OA induction, mice received intra-articular injections of the EZH2 inhibitor EPZ-6438 (10 µM, 10 µL) or vehicle into the right knee joint. At day 56, mice were euthanized and histological analyses were performed. Knee sections were stained with Safranin O and counterstained with Fast Green, and representative images were acquired using light microscopy (b). Cartilage degradation was evaluated using OARSI and Mankin scores (c), and synovial thickness was quantified (d). Data are presented as mean ± SEM (n=4 mice/group). Functional pain was assessed weekly throughout the experiment using von Frey filaments (e) and at day 56 using static weight-bearing distribution (f). Data are presented as median ± interquartile range (n=8 mice/group). Statistical significance is indicated as *p < 0.05 and p < 0.01. For longitudinal von Frey analysis (e), the vehicle-treated OA group was compared with the sham group (##) and the EPZ-6438-treated OA group (**).

Histological analyses revealed preservation of articular cartilage integrity following EZH2 inhibition, as demonstrated by reduced cartilage degradation scores compared with vehicle-treated animals (Figure 1b-c). In parallel, EPZ-6438 administration decreased synovial thickening, suggesting reduced synovial inflammatory remodeling (Figure 1d).

Importantly, EZH2 inhibition also improved functional outcomes. EPZ-6438-treated mice exhibited reduced mechanical hypersensitivity and improved weight distribution on the affected limb compared with vehicle-treated animals (Figure 1e-f).

Together, these results demonstrate that local pharmacological inhibition of EZH2 limits multiple features of MIA-induced joint pathology and associated pain-related behaviors, supporting a broader role of EZH2 beyond cartilage degeneration.

### EZH2 drives inflammatory and catabolic responses in OA synoviocytes

Because synovial inflammation contributes substantially to OA progression and pain sensitization, we next investigated whether synovial fibroblasts represent a cellular target of EZH2 activity. Human OA-derived fibroblast-like synoviocytes (FLS) were stimulated with IL-1β to reproduce an inflammatory OA-like environment.

IL-1β stimulation increased EZH2 expression in FLS, indicating that inflammatory signaling activates this epigenetic regulator in synovial cells (Figure 2a). Consistently, IL-1β increased the proportion of H3K27me3-positive cells, whereas EPZ-6438 treatment reduced H3K27me3 accumulation, confirming inhibition of EZH2 methyltransferase activity (Figure 2b).

**Figure 2:**
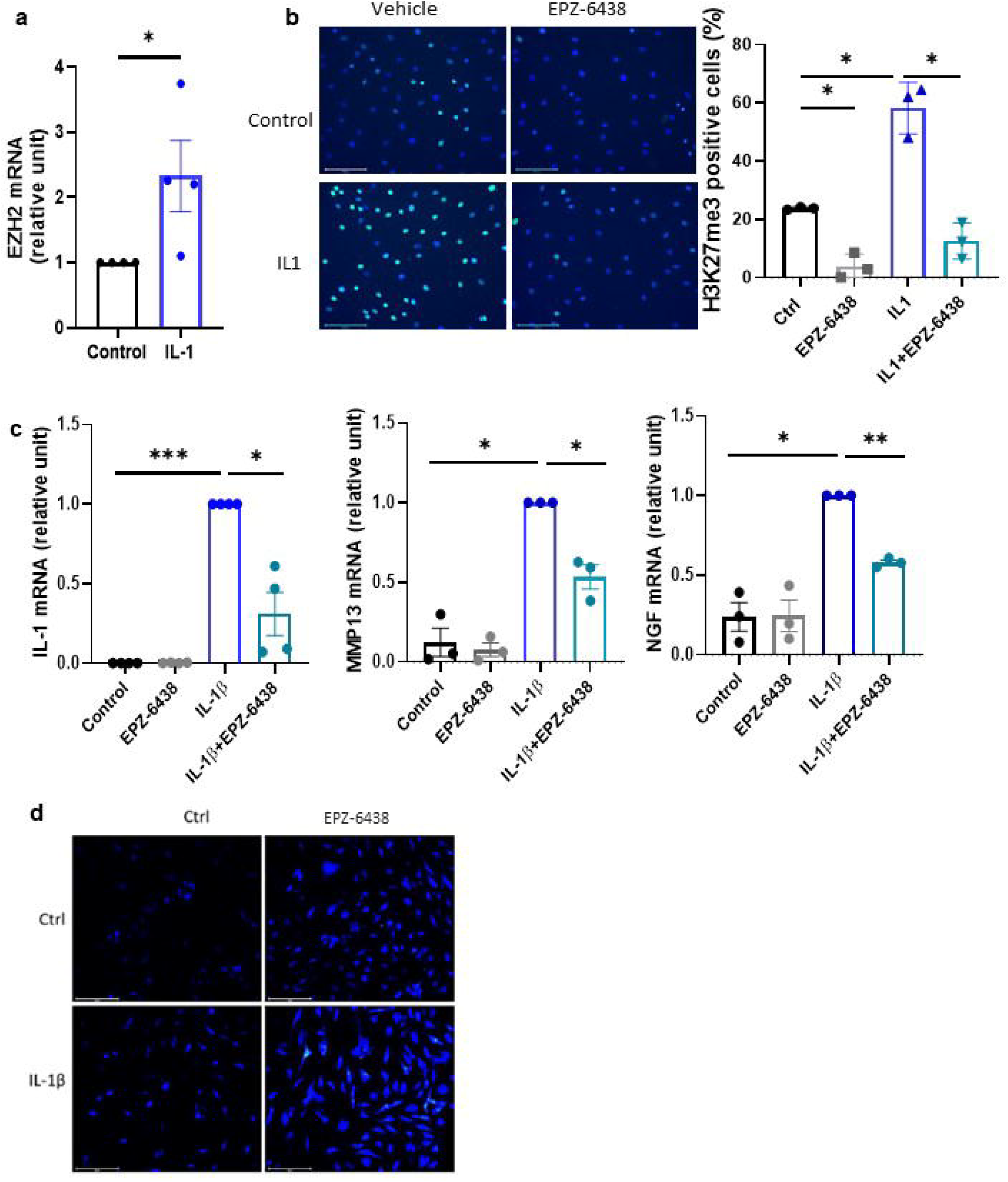
EZH2 inhibition modulates inflammatory, catabolic, pain-related and autophagic responses in fibroblast-like synoviocytes. Human fibroblast-like synoviocytes were stimulated with IL-1β (1 ng/mL) for 48 h. EZH2 mRNA expression was analyzed by RT-qPCR (a). Cells stimulated with IL-1β in the presence or absence of EPZ-6438 (10 µM) were stained for H3K27me3 (FITC, green) by immunocytochemistry, and nuclei were counterstained with DAPI. Representative images from three independent experiments are shown. Quantification of H3K27me3-positive cells is presented (b). The expression of inflammation-, catabolism- and pain-related genes (IL1B, MMP13 and NGF) was analyzed by RT-qPCR (c). Data are presented as mean ± SEM from at least three independent biological experiments. Statistical significance is indicated as *p < 0.05, **p < 0.01 and ***p < 0.001. Autophagy was assessed using a fluorescent autophagosome detection assay. Representative images from three independent experiments are shown (d).

Functionally, EZH2 inhibition markedly altered the inflammatory phenotype of activated synoviocytes. EPZ-6438 reduced IL-1β-induced expression of inflammatory, catabolic, and pain-associated mediators, including IL-1β, MMP13, and NGF (Figure 2c). In addition, EPZ-6438 increased autophagosome accumulation in FLS (Figure 2d), suggesting restoration of cellular homeostatic mechanisms.

Together, these findings identify synoviocytes as a major cellular target of EZH2 activity and suggest that EZH2 coordinates inflammatory and pain-associated responses within the synovial compartment.

### Proteomic profiling identifies inflammatory, metabolic and neuronal-associated pathways regulated by EZH2 inhibition

To characterize the molecular consequences of EZH2 inhibition in inflammatory synoviocytes, we performed quantitative proteomic profiling of IL-1β-stimulated OA-derived FLS treated with EPZ-6438.

EZH2 inhibition induced significant changes in the abundance of 128 proteins (p < 0.05; fold change >1.2), with most altered proteins showing increased abundance following treatment (Figure 3a; Table S1). Functional enrichment analysis revealed that EZH2-dependent proteins were associated with pathways relevant to OA biology, including inflammatory signaling, extracellular matrix regulation, and autophagy-related processes (Figure 3b).

**Figure 3:**
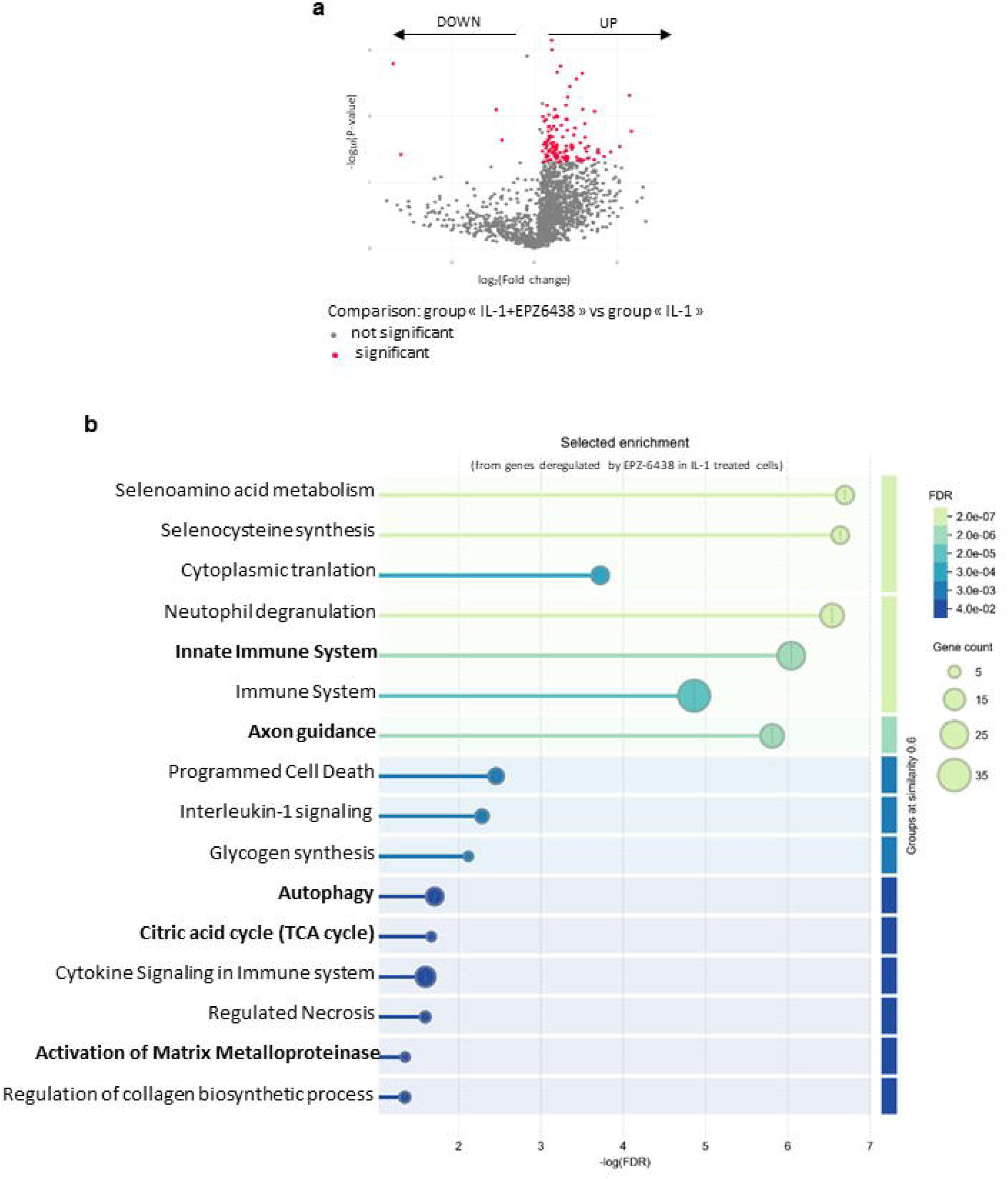
Proteomic profiling identifies inflammatory, metabolic and neuronal-associated pathways regulated by EZH2 inhibition. Human fibroblast-like synoviocytes from OA patients (n=3 independent biological replicates) were stimulated with IL-1β (1 ng/mL) for 48 h in the presence or absence of EPZ-6438 (10 µM). Total proteins were extracted and analyzed by label-free quantitative proteomics. Differentially abundant proteins between IL-1β and IL-1β + EPZ-6438 conditions were identified using a volcano plot representation (a). Proteins displaying a fold change ≥1.2 and a p-value <0.05 were considered differentially abundant. Gene Ontology Biological Process and Reactome pathway enrichment analyses were performed using STRING database (b).

Unexpectedly, enrichment analysis also revealed a strong association with neuronal-associated biological processes, particularly axon guidance pathways. These findings suggest that EZH2 influences molecular pathways associated with neuroimmune communication in inflammatory synoviocytes.

Several metabolic pathways were also enriched among EZH2-regulated proteins, including the TCA cycle and glycogen metabolism.

### EZH2 inhibition modulates representative metabolic and neuronal genes in inflammatory synoviocytes

Because proteomic analysis identified several metabolism-related pathways, we first examined LDHA expression, which was significantly reduced by EPZ-6438 (Figure 4a). However, Seahorse analysis did not reveal significant alterations in glycolytic or mitochondrial function (Figure S1), suggesting that EZH2 inhibition modifies metabolic gene expression without inducing major bioenergetic remodeling. We next investigated the neuronal guidance molecule NTN1, which was selected because of its reported role in sensory nerve ingrowth and OA pain. IL-1β increased NTN1 expression, whereas EPZ-6438 markedly reduced this induction (Figure 4b). These data suggest that EZH2 contributes to inflammatory activation of neuro-associated signaling pathways in synoviocytes.

**Figure 4:**
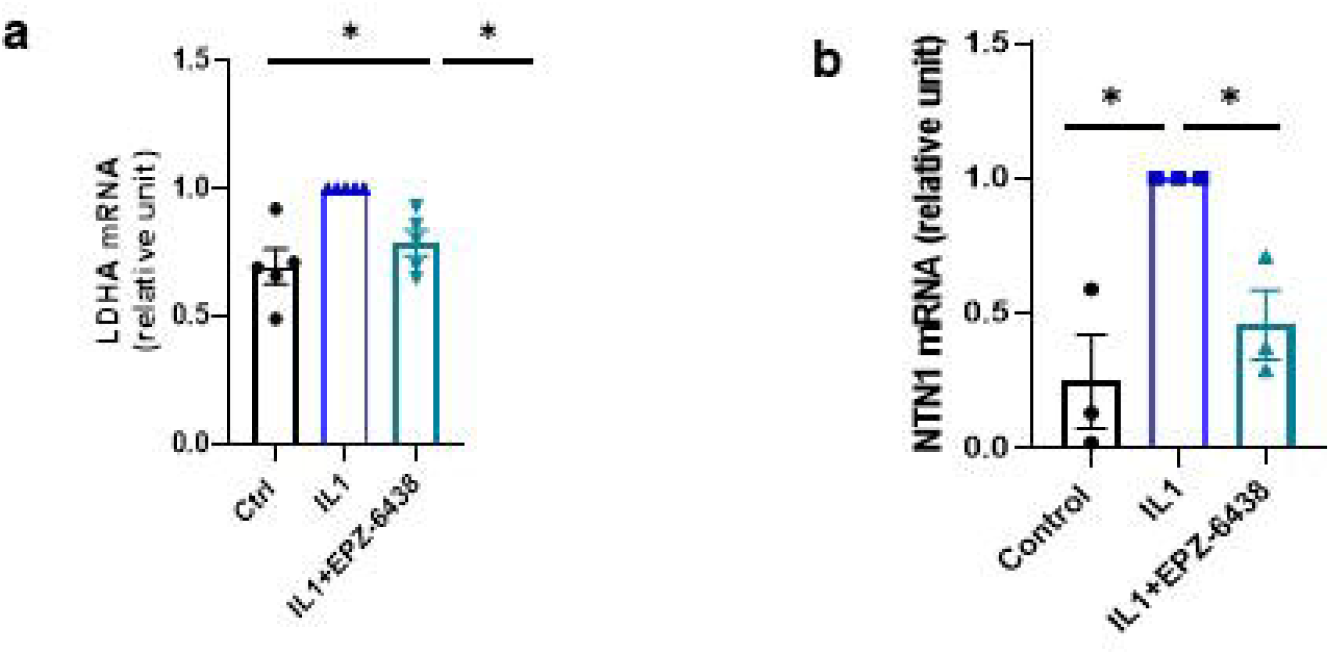
EZH2 inhibition modulates representative metabolic and neuronal genes in inflammatory synoviocytes. Human fibroblast-like synoviocytes were stimulated with IL-1β (1 ng/mL) for 48 h in the presence or absence of EPZ-6438 (10 µM). RNA was extracted and LDHA (a) and NTN1 (b) mRNA expression was analyzed by RT-qPCR. Data are presented as mean ± SEM from at least three independent OA patients. Statistical significance is indicated as *p < 0.05.

### ChIP-seq identifies neurodevelopmental genes as potential EZH2 targets in inflammatory synoviocytes

To further investigate potential epigenetic mechanisms underlying these responses, genome-wide EZH2 ChIP-seq analysis was performed in IL-1β-stimulated FLS. Inflammatory stimulation induced the recruitment of EZH2 to promoter regions corresponding to 285 genes. Functional enrichment analysis revealed a significant association with nervous system development and axon guidance pathways, including genes involved in neuronal differentiation and developmental regulation (Figures 5a-b and S2).

**Figure 5:**
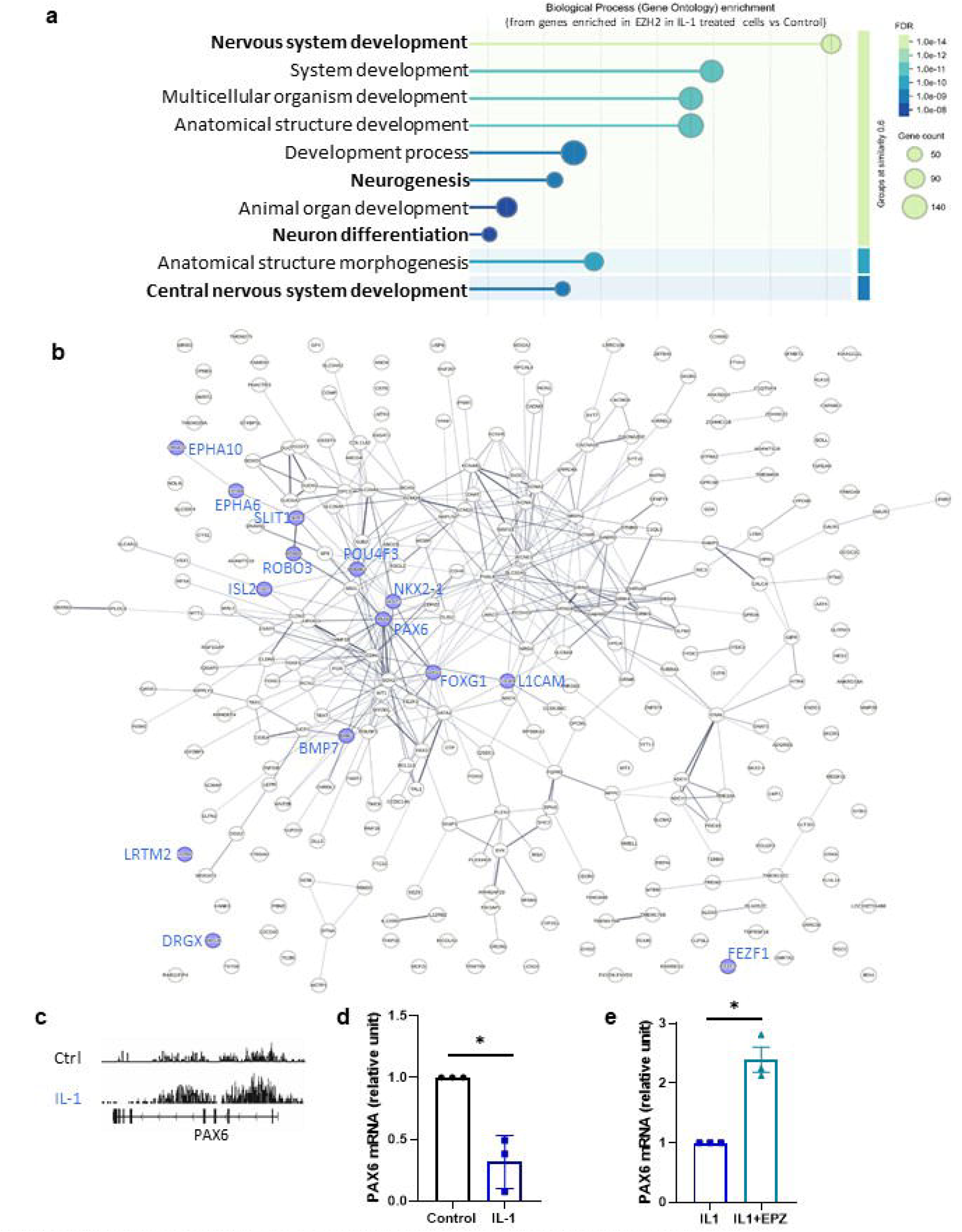
ChIP-seq identifies neurodevelopmental genes as potential EZH2 targets in inflammatory synoviocytes. Fibroblast-like synoviocytes were incubated for 48 h with IL-1β (1 ng/ml), and EZH2 genomic occupancy was investigated by ChIP-seq using an EZH2-specific antibody. Sequencing data were analyzed by identifying common and condition-specific EZH2 peaks. Functional enrichment analysis (Gene Ontology Biological Processes) was performed on genes associated with EZH2 promoter peaks specifically identified under IL-1β stimulation using STRING database (a). The protein-protein interaction network of genes associated with IL-1β-specific EZH2 peaks is shown, with genes related to the “axon guidance” pathway highlighted in blue (b). Representative ChIP-seq peaks corresponding to the PAX6 locus are shown (c). PAX6 mRNA expression was analyzed by RT-PCR (d), and the effect of EPZ-6438 treatment on PAX6 expression was assessed (e). ChIP-seq experiments were performed using biological replicates from two independent OA patients. RT-PCR experiments were independently performed using biological replicates from three different OA patients Data are presented as mean ± SEM. *: p-value < 0.05.

Among these candidates, PAX6 was selected for further validation because of its established role in neuronal development and axonal guidance. EZH2 enrichment was detected at the PAX6 locus (Figure 5c), and inflammatory stimulation reduced PAX6 expression, whereas EZH2 inhibition restored PAX6 expression levels (Figure 5d-e).

Together, these data identify neurodevelopmental genes as candidate EZH2 targets in inflammatory synoviocytes, providing candidate molecular links between epigenetic regulation and neuronal-associated pathways in OA.

### EZH2 inhibition reduces inflammatory macrophage activation

Because macrophages are major regulators of synovial inflammation and OA-associated pain, we next examined whether EZH2 also controls macrophage activation.

THP-1-derived macrophages polarized toward an inflammatory M1-like phenotype in the presence of EPZ-6438 displayed reduced expression of inflammatory mediators, including IL-1β and CXCL1 (Figure 6a). Similar effects were observed in macrophages differentiated from OA patient-derived bone marrow cells, where EZH2 inhibition decreased IL-1β, IL-6, and CXCL1 expression (Figure 6b).

**Figure 6:**
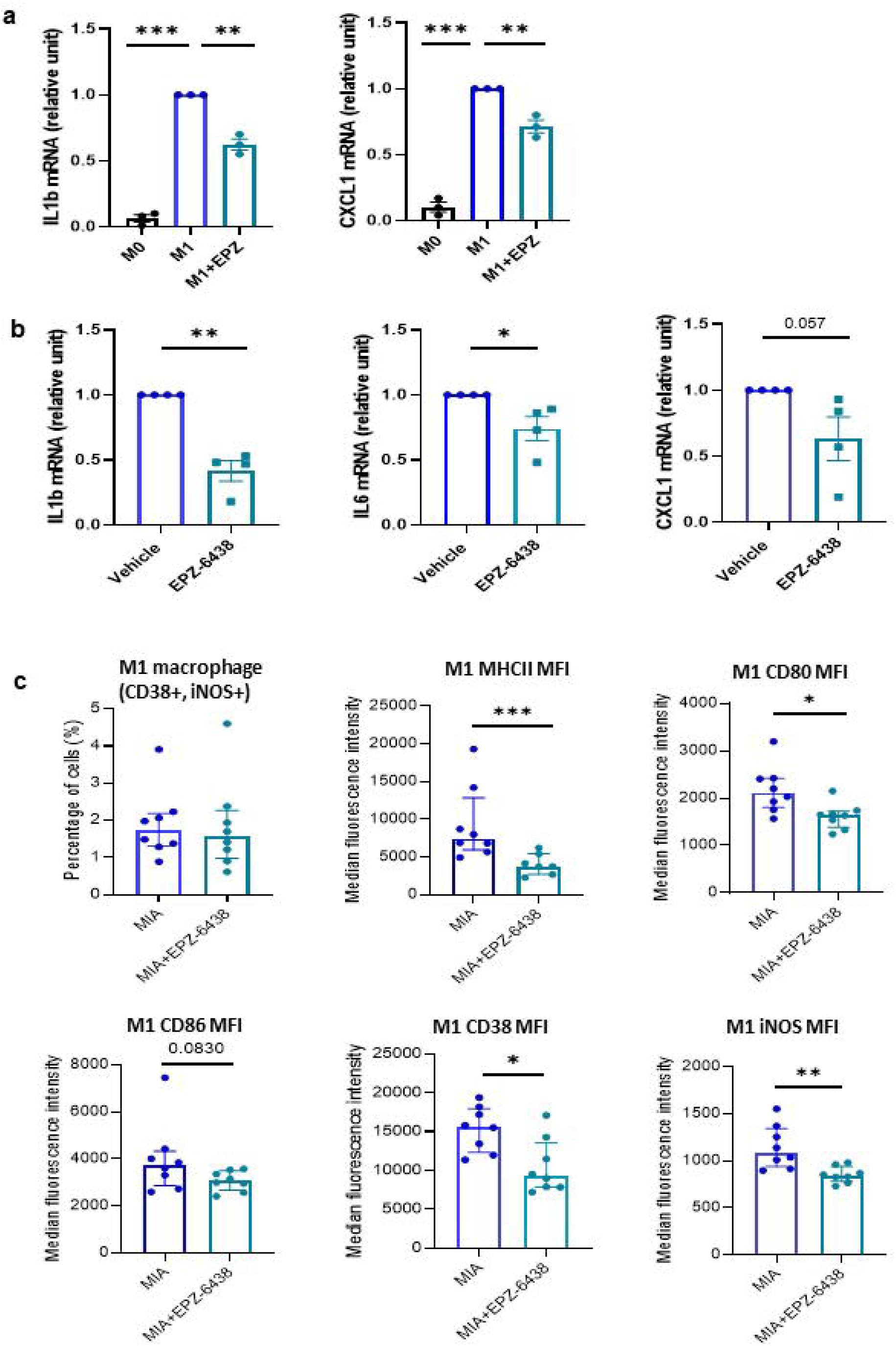
EZH2 inhibition reduced the inflammatory phenotype of macrophages during OA. Macrophage differentiation and M1-like polarization were induced from the THP-1 monocytic cell line (a) or from human bone marrow-derived cells isolated from OA patients (b) in the presence or absence of EPZ-6438. The expression of pro-inflammatory markers (IL-1β, IL-6 and CXCL1) was analyzed by RT-qPCR. Experiments using THP-1 cells were independently repeated at least three times. Experiments using human bone marrow-derived cells were performed using biological replicates from independent OA patients. Data are presented as mean ± SEM. *: p-value < 0.05; **: p-value < 0.01; ***: p-value < 0.001. Spleens from OA mice described in Figure 1 and treated with vehicle or EPZ-6438 were collected and analyzed by flow cytometry. Cells were stained for macrophage-associated markers (c). Samples were acquired using a FACSVerse flow cytometer and analyzed with FlowJo 7.6.5 software. Results are presented as median ± interquartile range (n=8 mice). *: p-value < 0.05; **: p-value < 0.01.

To determine whether EZH2 inhibition influenced immune responses *in vivo*, immune cells from OA mice treated with EPZ-6438 were analyzed by flow cytometry. EPZ-6438-treated animals displayed reduced expression of inflammatory macrophage-associated markers, including MHCII, CD80, CD38, and iNOS (Figures 6c and S3).

These results demonstrate that EZH2 contributes to macrophage inflammatory activation and suggest that modulation of immune responses may participate in the protective effects of EZH2 inhibition during OA.

### EZH2 inhibition suppresses osteoclast differentiation

Subchondral bone remodeling represents another important component of OA progression and pain. Since EZH2 has previously been implicated in osteoclast differentiation, we investigated whether EZH2 inhibition affects osteoclastogenesis in OA-derived cells.

Bone marrow-derived progenitors isolated from OA patients were induced to differentiate into osteoclasts in the presence or absence of EPZ-6438. EZH2 inhibition significantly reduced formation of TRAP-positive multinucleated osteoclasts (Figure 7a-b).

**Figure 7:**
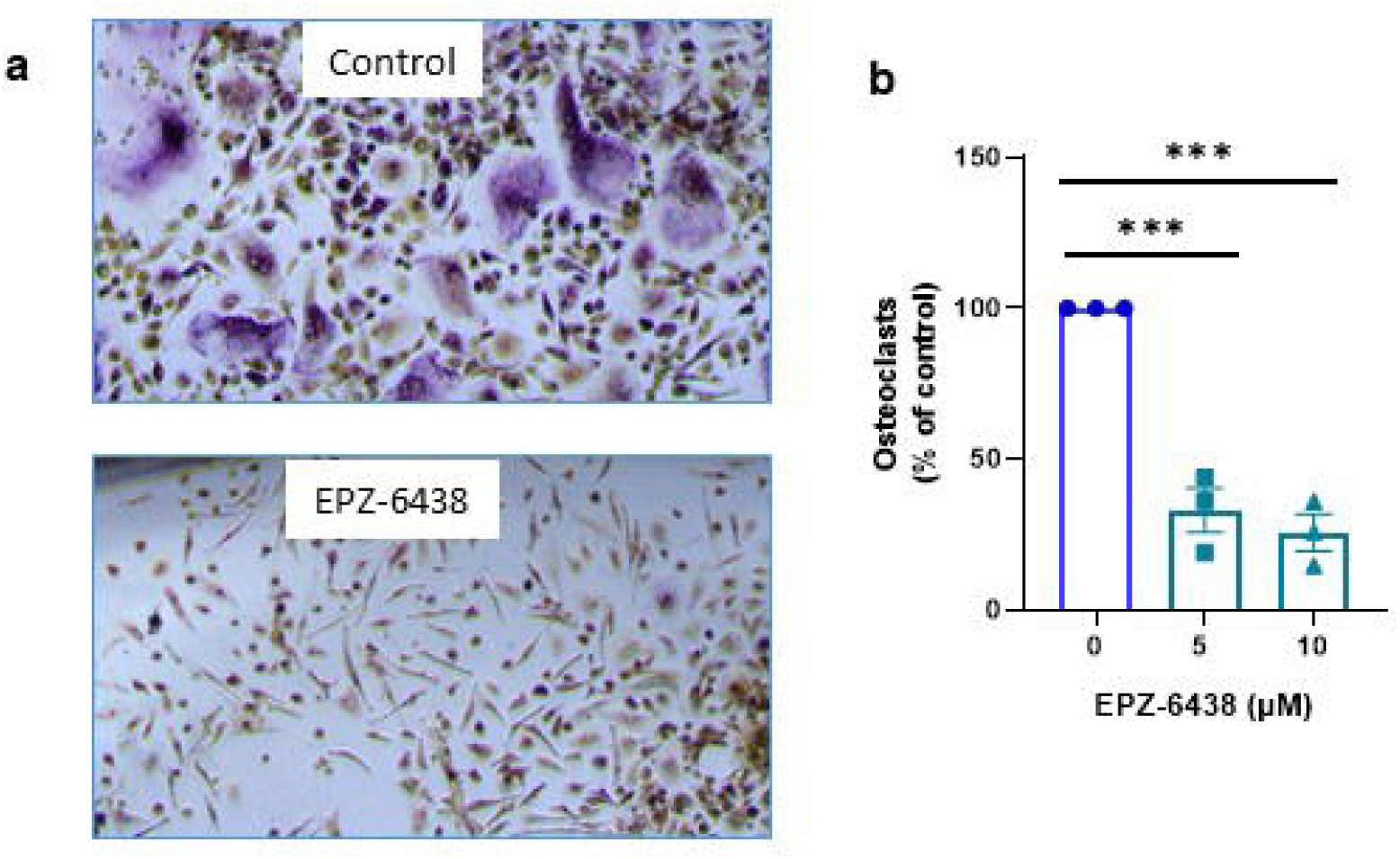
EZH2 inhibition reduced osteoclastogenesis. Bone marrow-derived cells were isolated from OA patients. After ACK lysis, cells were cultured at 300,000 cells/cm^2^ in α-MEM supplemented with 10% FBS, penicillin-streptomycin, 50 µM β-mercaptoethanol and osteoclastogenic cytokines (RANK-L, 30 ng/mL; M-CSF, 25 ng/mL) in the presence or absence of EPZ-6438 (5 or 10 µM). When multinucleated giant cells were observed (approximately seven days), osteoclast differentiation was assessed by TRAP staining according to the manufacturer’s protocol. Representative images are shown (a), and TRAP-positive multinucleated osteoclasts were counted and expressed as a percentage of the untreated control condition (set at 100%) (b). Results are presented as mean ± SEM from three independent OA patients (n=3). ***: p-value < 0.001.

These findings indicate that EZH2 contributes to osteoclastogenic responses in OA-derived cells and further extend its potential role beyond cartilage and synovial inflammation to additional joint remodeling processes.

## Discussion

Osteoarthritis is increasingly recognized as a whole-joint disease in which reciprocal interactions between cartilage, synovium, immune cells, subchondral bone and sensory nerves collectively drive structural damage and chronic pain. This evolving concept challenges the traditional cartilage-centered view of OA and highlights the need to identify upstream regulators capable of coordinating pathogenic responses across multiple joint compartments. In the present study, we demonstrate that pharmacological inhibition of EZH2 attenuates structural joint damage and pain-related behaviors while simultaneously modulating inflammatory, neuronal-associated and osteoimmune pathways in several OA-relevant cell populations. Together, our findings extend the role of EZH2 beyond cartilage degeneration and support a broader role for EZH2 in coordinating multiple OA-relevant cellular responses.

Previous studies, including our own, have demonstrated that EZH2 promotes cartilage degeneration by enhancing chondrocyte hypertrophy, inflammatory responses and extracellular matrix degradation (6–9). Our data considerably broaden this view by showing that EZH2 activity is not restricted to cartilage but is also engaged in synovial fibroblasts, inflammatory macrophages and osteoclast precursors. This broader activity is consistent with the emerging concept that epigenetic regulators function as integrators of environmental signals, enabling coordinated transcriptional responses to inflammatory, mechanical and metabolic stress. Rather than acting within a single tissue, EZH2 therefore appears to orchestrate pathological responses across several cellular compartments that collectively contribute to OA progression.

A major finding of this study is the identification of fibroblast-like synoviocytes as an important EZH2-responsive compartment. Synovial fibroblasts are increasingly recognized as active drivers of OA progression through the production of inflammatory cytokines, matrix-degrading enzymes and soluble mediators capable of influencing immune cell recruitment and nociceptive signaling. In our study, inflammatory stimulation increased both EZH2 expression and H3K27me3 accumulation, while pharmacological inhibition reduced IL1B, MMP13 and NGF expression and enhanced autophagy. These observations indicate that EZH2 contributes to the establishment of a pathogenic synovial phenotype and suggest that restoration of cellular homeostasis through enhanced autophagy may participate in the protective effects of EZH2 inhibition. Together with recent transcriptomic studies demonstrating extensive inflammatory and stromal remodeling within OA synovium (15), our findings reinforce the concept that synovial fibroblasts constitute a major therapeutic target for epigenetic intervention.

One of the most intriguing observations of this work is the identification of neuronal-associated pathways among the principal molecular programs regulated by EZH2. Although pain is the predominant clinical manifestation of OA, the molecular mechanisms linking synovial inflammation to sensory nerve remodeling remain incompletely understood. Our proteomic analyses unexpectedly identified axon guidance among the most significantly enriched pathways following EZH2 inhibition. Consistent with these observations, EPZ-6438 reduced inflammatory induction of NTN1, a guidance molecule previously implicated in sensory nerve sprouting and pain sensitization during OA (18,19). Furthermore, genome-wide ChIP-seq identified inflammation-dependent recruitment of EZH2 to promoters of genes involved in nervous system development, including PAX6, whose expression was restored after EZH2 inhibition. Although these findings do not establish a direct causal role for PAX6 in OA pathogenesis, they identify neurodevelopmental genes as previously unrecognized EZH2-associated targets in inflammatory synoviocytes. More broadly, they suggest that epigenetic regulation within stromal cells may contribute to neuroimmune communication in the osteoarthritic joint, thereby providing one potential explanation for the reduction in pain-related behaviors observed after EZH2 inhibition.

Beyond resident synovial cells, our study demonstrates that EZH2 also regulates inflammatory and remodeling responses in immune and skeletal compartments. EPZ-6438 reduced inflammatory activation of macrophages both in vitro and in vivo, consistent with previous studies identifying EZH2 as a regulator of macrophage polarization (13,20). Because activated macrophages contribute to synovitis, cytokine production and peripheral sensitization (12), modulation of macrophage phenotype is likely to participate in the overall therapeutic effects observed in vivo. Likewise, inhibition of osteoclast differentiation extends previous reports describing EZH2 as a positive regulator of osteoclastogenesis (14,21). Since subchondral osteoclast activity promotes bone remodeling, angiogenesis and sensory nerve ingrowth during OA progression (22), these findings are consistent with a role of EZH2 in regulating osteoimmune mechanisms contributing both to structural damage and pain generation.

Interestingly, quantitative proteomics also identified several metabolic pathways among proteins regulated by EZH2 inhibition. However, despite reduced expression of LDHA, complementary Seahorse analyses did not reveal major alterations in glycolytic or mitochondrial activity. These results suggest that the metabolic signatures identified by proteomics more likely reflect secondary adaptations accompanying the anti-inflammatory response than a primary metabolic reprogramming induced by EZH2 inhibition. Future studies combining metabolic flux analyses with longitudinal kinetic assessments will be important to define the temporal dynamics of metabolic responses to EZH2 inhibition.

Several limitations should nevertheless be acknowledged. First, the present study relies primarily on pharmacological inhibition of EZH2 using EPZ-6438. Although this clinically approved inhibitor offers strong translational relevance, cell-specific genetic approaches will be required to determine the relative contribution of individual joint compartments to the therapeutic effects observed in vivo. Second, while the MIA model faithfully reproduces pain-related alterations and several pathological features of OA, it does not encompass the full complexity of human disease. Confirmation of these findings in additional experimental models will therefore be important. Finally, although our study identifies candidate neuroimmune pathways involving NTN1 and PAX6, their precise functional contribution to sensory nerve remodeling and OA pain remains to be experimentally demonstrated.

In conclusion, our findings identify EZH2 as a central epigenetic regulator associated with inflammatory, neuroimmune and osteoimmune pathways in OA. Rather than acting exclusively on cartilage, EZH2 appears to integrate pathogenic mechanisms operating within stromal, immune and skeletal compartments, ultimately influencing both structural progression and pain. These results provide a conceptual framework supporting local EZH2 inhibition as a promising strategy to simultaneously target multiple disease-driving mechanisms in osteoarthritis.

## Material and methods

### Reagents

The EZH2 inhibitor EPZ-6438 (tazemetostat; Selleckchem, Souffelweyersheim, France) was dissolved in dimethyl sulfoxide (DMSO; Dutscher, Bernolsheim, France) at a stock concentration of 10 mM and stored at -20°C until use. For *in vitro* experiments, EPZ-6438 was diluted directly into culture medium to obtain a final concentration of 10 µM. For *in vivo* experiments, EPZ-6438 was diluted in sterile saline solution to obtain a final concentration of 10 µM for intra-articular injection. Human recombinant IL-1β (SRP3083, Sigma-Aldrich, St Louis, USA) was reconstituted in DPBS supplemented with 0.1% BSA at 1 µg/mL and used at a final concentration of 1 ng/mL. Phorbol 12-myristate 13-acetate (PMA; MedChemExpress, Monmouth Junction, USA) was dissolved in DMSO at a stock concentration of 20 mM. Recombinant human interferon gamma (IFNγ; PreproTech, Cranbury, USA) was reconstituted in DPBS containing 0.1% BSA at 10 µg/mL.

### Animals

Twenty-seven 8-week-old male C57BL/6J mice (Janvier Labs, France) were used in this study. Male mice were selected as they represent the most commonly used sex in monoiodoacetate (MIA)-induced osteoarthritis models and allow comparison with previous studies investigating EZH2 inhibition in experimental OA. All animals were housed under controlled conditions (23 ± 2°C, 12-hour reversed light/dark cycle) at the Centre Universitaire de Ressources Biologiques (CURB, University of Caen Normandy, France), with free access to food and water. Animal procedures were performed between 8:00 AM and 5:00 PM under dim illumination (6 lux).

All experimental procedures complied with the European Directive 2010/63/EU and French legislation on animal experimentation (decree 87-848). The study was conducted at the GIP CYCERON/CURB facility (approval #D14118001) and was approved by the French Ministry of Education and Research and the regional ethics committee CENOMEXA (APAFIS number #16185). Experimental procedures were performed according to ARRIVE guidelines. Every effort was made to minimize animal suffering and reduce the number of animals used.

Animals were randomly allocated to the experimental groups. A subset of animals was used for behavioral and histological analyses, whereas spleens and lymph nodes from treated animals were collected for flow cytometry analyses.

### Induction of experimental osteoarthritis and treatment

The experiments were performed according the timeline presented in figure 1A.

Experimental osteoarthritis was induced by intra-articular injection of monosodium iodoacetate (MIA; Sigma-Aldrich) as previously described (23). Briefly, mice received an intra-articular injection of 0.75 mg of MIA diluted in 10 µL of sterile saline solution into the right knee joint at day 0. Sham animals received an intra-articular injection of vehicle (10 µL sterile saline solution) at the same time point. EPZ-6438 treatment was administered intra-articularly into the right knee joint at days 2, 7, and 30 after OA induction. EPZ-6438 was diluted from a 10 mM stock solution prepared in DMSO to a final concentration of 10 µM in sterile saline solution (10 µL/injection, corresponding to 0.043 µg EPZ-6438 per injection). Control OA animals received intra-articular injections of vehicle at the same time points.

Animals were euthanized at day 56 after OA induction by cervical dislocation under deep anesthesia (5% isoflurane, 70% N_2_O/30% O_2_).

### Histological analysis and cartilage scoring

At day 56, knee joints were collected and fixed in 10% formalin (4% formaldehyde) for 48 hours. Samples were subsequently decalcified in neutral 10% EDTA/PBS for 5 days, cryoprotected in 30% sucrose solution for 48 hours, embedded in OCT compound, and stored at −80°C.

Sections (10 µm thickness) were obtained using a cryostat (CM3050S, Leica Biosystems, Nanterre, France), mounted on Superfrost Plus slides (Thermo Fisher Scientific), and stored at −80°C until staining. Sections were stained with Safranin O and counterstained with Fast Green as previously described (23). Cartilage degradation was assessed using the OARSI scoring system adapted for murine osteoarthritis (24) and the Mankin score (25). Synovial alterations were evaluated by measuring synovial membrane thickness. Images were acquired using a Leica DM6 microscope equipped with a DMC2900 camera.

### Functional pain assessment

OA-associated pain behavior was assessed longitudinally using von Frey filament testing and at the end of the experiment using static weight-bearing (SWB) analysis.

Before behavioral experiments, mice underwent a three-week habituation period with regular handling twice weekly to reduce stress associated with manipulation. Behavioral procedures were performed at the HANDIFORM platform (CYCERON/CURB, Caen, France).

Mechanical tactile sensitivity was evaluated once a week after OA induction using von Frey filaments (Ugo Basile, Salon-de-Provence, France), as previously described (23). Mice were placed individually in elevated grid cages and allowed to acclimatize for 1 hour before testing. Mechanical stimulation was applied to the plantar surface of each hind paw using calibrated nylon filaments. A positive response was defined as paw withdrawal, snapping, or licking following stimulation. Three positive responses among a maximum of five stimulations were considered indicative of a response threshold. Results were expressed as the ratio between withdrawal responses of the OA-injected right paw and the contralateral left paw.

Static weight-bearing distribution was assessed once at day 56 using a Bioseb SWB system (Vitrolles, France). After one day of habituation to the apparatus, ten consecutive measurements were performed for each animal. The mean weight supported by the right hind paw was expressed relative to the total weight supported by both hind paws (right/(right + left)).

### Fibroblast-like synoviocyte cultures and treatment

Human fibroblast-like synoviocytes (FLS) were isolated from synovial membrane samples obtained from patients with osteoarthritis undergoing total hip replacement surgery (mean age: 75 years). Both male and female patients were included in the study. The experimental protocol was approved by the local ethics committee (“Comité de Protection des Personnes Nord-Ouest III”, agreement #A13-D46-VOL.19). All participants provided written informed consent prior to surgery, and all procedures were conducted in accordance with applicable ethical guidelines and regulations.

Synovial membrane samples were enzymatically digested with collagenase type I (2 mg/mL, 12 h; Thermo Fisher Scientific, Waltham, USA) as previously described (26). Released cells were cultured in high-glucose Dulbecco’s Modified Eagle Medium (DMEM) supplemented with glutamine and sodium pyruvate (Dutscher, Bernolsheim, France), 10% fetal bovine serum (FBS; Dutscher), and penicillin-streptomycin (Lonza, Basel, Switzerland), under standard conditions (37°C, humidified atmosphere containing 5% CO_2_). Cells were routinely tested for mycoplasma contamination by PCR.

For experiments, FLS between passages 3 and 8 were seeded at approximately 1 × 10^4^ cells/cm^2^. At confluence (approximately 5 days after seeding), cells were stimulated with recombinant human IL-1β (1 ng/mL) in the presence or absence of EPZ-6438 (10 µM) for the indicated duration.

### Immunocytochemistry and H3K27me3 staining

H3K27me3 staining was performed as previously described (7). Briefly, cells were fixed and incubated overnight with an anti-H3K27me3 antibody (C36B11, Cell Signaling Technology). After washing, cells were incubated with fluorescent secondary antibody (anti-rabbit IgG-FITC, sc-2012, Santa Cruz Biotechnology) and counterstained with DAPI. Fluorescence images were acquired using an EVOS FL Auto 2 Cell Imaging System (Thermo Fisher Scientific). H3K27me3 staining intensity was quantified from independent biological replicates.

### mRNA gene expression analysis

Total RNA was extracted from cells using the RNeasy Mini Kit (Qiagen, Hilden, Germany) according to the manufacturer’s instructions. A DNase treatment step (AMPD1, Sigma-Aldrich) was performed to remove genomic DNA contamination. Reverse transcription was performed using 1 µg of total RNA and the M-MLV reverse transcriptase kit (Invitrogen, Carlsbad, USA). Quantitative real-time PCR was performed using the StepOne Plus Real-Time PCR System (Applied Biosystems, Courtaboeuf, France). PCR reactions were performed using Power SYBR Green Master Mix (Applied Biosystems) with gene-specific primers (200 nM each). Relative gene expression levels were calculated using the comparative 2^-ΔΔCt^ method and normalized to the housekeeping gene RPL13.

### Autophagy assessment

Autophagy was evaluated using the Autophagy Assay Kit (MAK138, Sigma-Aldrich), which detects autophagosome formation through a fluorescent autophagosome marker (λex = 333 nm; λem = 518 nm). FLS were seeded at 10,000 cells/well in 96-well plates and treated for 48 h with IL-1β (1 ng/mL) in the presence or absence of EPZ-6438 (10 µM). Cells were then incubated with the autophagy detection reagent for 30 min at 37°C, washed according to the manufacturer’s instructions, and fluorescence intensity was measured at excitation/emission wavelengths of 360/520 nm.

### Proteomic experiments

Fibroblast-like synoviocytes derived from three independent osteoarthritis patients were cultured and treated with IL-1β (1 ng/mL) in the presence or absence of EPZ-6438 (10 µM). Cells were lysed and proteins were extracted using radioimmunoprecipitation assay (RIPA) buffer (50 mM Tris-HCl pH 7.5, 1% Igepal CA-630, 150 mM NaCl, 1 mM EGTA, 1 mM NaF, and 0.25% sodium deoxycholate) supplemented with protease and phosphatase inhibitors, as previously described (26). Protein concentrations were determined using the Bradford assay (Bio-Rad, Marnes-la-Coquette, France). Samples were then submitted to the PROTEOGEN proteomics platform at the University of Caen Normandy for mass spectrometry analysis.

Proteomic sample preparation and mass spectrometry acquisition were performed as previously described (26). The mass spectrometer was operated in parallel accumulation–serial fragmentation (PASEF) mode. Ten PASEF MS/MS scans were acquired within a 1.25 s cycle time, with precursor ion charge states ranging from 2 to 5.

### Identification of differentially abundant proteins and enrichment analysis

Relative protein abundance was quantified using the label-free quantification (LFQ) workflow implemented in PEAKS X+ software. Feature detection was performed independently for each sample using an expectation-maximization algorithm. Peptide features were aligned across replicates using retention time alignment algorithms. Mass error tolerance was set at 20 ppm, ion mobility tolerance at 0.07 (1/k0), and retention time tolerance at 10 min. Sample normalization was performed based on total ion current (TIC). Protein abundance was calculated using the summed intensity of the three most abundant unique peptides.

Differentially abundant proteins between experimental conditions were identified using paired Student’s t-test implemented in Perseus. Proteins displaying a fold change ≥1.2 and a p-value <0.05 were considered significantly deregulated. Volcano plots were generated using Genoppi. Gene Ontology biological processes and Reactome pathway enrichment analyses were performed using STRING database version 12.0 (27). Proteomic data are publicly available through the iProX integrated proteome resources database (dataset ID: IPX0011070000).

### ChiP-Seq

ChIP-seq experiments were performed by the Diagenode ChIP-seq/ChIP-qPCR Profiling Service using an anti-EZH2 antibody. ChIP-seq libraries were prepared and sequenced in paired-end mode (2 × 150 bp) on an Illumina NovaSeq 6000 platform using NovaSeq Control Software version 1.6.0. Sequencing quality was assessed using FastQC. Reads were aligned to the human reference genome (GRCh38/hg38) using BWA. Reads mapping to ENCODE blacklisted regions, multimapping reads, and PCR duplicates were removed using samtools. Alignment files were converted to BED format using BEDTools v2.17, and EZH2-enriched regions (peaks) were identified using MACS2. Peaks were visualized using the Integrative Genomics Viewer (IGV). Differential EZH2 binding profiles between experimental conditions were analyzed by identifying common and condition-specific peaks. Genes associated with condition-specific EZH2 promoter binding sites under IL-1β stimulation were subjected to Gene Ontology Biological Process enrichment analysis using STRING database (version 12.0). ChIP-seq data have been deposited in the Gene Expression Omnibus database under accession number GSE300508.

### Macrophage differentiation and polarization

Macrophage differentiation and polarization of the THP-1 monocytic cell line (Merck, Sigma-Aldrich) were performed according to a protocol adapted from Baxter et al. (28). THP-1 cells were maintained in RPMI medium supplemented with 10% fetal bovine serum and penicillin-streptomycin at 37°C in a humidified atmosphere containing 5% CO_2_. Cells were routinely tested for mycoplasma contamination by PCR.

For macrophage differentiation, THP-1 cells were seeded at 50 × 10^3^ cells/cm^2^ and stimulated with PMA (250 ng/mL) for 24 h. Cells were then washed and cultured for 72 h in PMA-free medium to obtain resting macrophages (M0-like macrophages). Pro-inflammatory polarization was induced by stimulation with IFNγ (20 ng/mL) and LPS from E. coli O55 (250 ng/mL) in the presence or absence of EPZ-6438 (10 µM).

Human bone marrow samples were collected from OA patients undergoing total hip replacement surgery (mean age = 70 years). The experimental protocol was approved by the local ethics committee, named “Comité de Protection des Personnes Nord Ouest III” (same agreement as for human synovium, #A13-D46-VOL.19). After red blood cell depletion using ACK lysing buffer, cells were cultured in DMEM supplemented with 10% fetal bovine serum, penicillin-streptomycin, β-mercaptoethanol (50 µM) for four days. Cells were subsequently stimulated with M-CSF (50 ng/mL) and LPS (50 ng/mL) in the presence or absence of EPZ-6438 (10 µM) for two days to induce a pro-inflammatory macrophage phenotype. Each experiment was performed using independent biological samples from different OA patients.

### Flow cytometry

Spleens and lymph nodes (brachial, cervical, and inguinal lymph nodes) were dissociated through 40 µm cell strainers (Falcon). Cells were resuspended in staining buffer, and Fc receptors were blocked with anti-CD16/32 antibodies (10 µg/mL; clone 2.4G2, 553142, BD Biosciences) for 15 min at 4°C. Cells were then stained with fluorochrome-conjugated monoclonal antibodies for surface markers for 10 min in the dark at 4°C. For intracellular staining, cells were fixed and permeabilized using the Inside Stain kit according to the manufacturer’s instructions (Miltenyi Biotec) before intracellular antibody staining.

The following antibodies were used for immune cell characterization: CD11b (REA592 VioBlue, 130-113-810, Miltenyi Biotec), F4/80 (REA126 PE-Vio770, 130-118-459, Miltenyi Biotec), CD80 (REA983 APC-Vio770, 130-116-463, Miltenyi Biotec), CD86 (REA1190 PE, 130-122-129, Miltenyi Biotec), MHC-II (REA813 FITC, 130-112-386, Miltenyi Biotec), CD38 (REA616 PerCP-Vio700, 130-109-260, Miltenyi Biotec), and iNOS (REA982 APC, 130-116-423, Miltenyi Biotec). Macrophages were identified based on CD11b and F4/80 expression, and inflammatory activation was assessed by the expression of CD80, CD86, MHC-II, CD38, and iNOS. Samples were acquired using a FACSVerse flow cytometer (BD Biosciences), and data were analyzed using FlowJo software version 7.6.5 (TreeStar Inc.).

### Osteoclastogenesis

Human bone marrow samples were collected from patients with osteoarthritis undergoing total hip replacement surgery (mean age⍰=⍰73⍰years). After red blood cell depletion using ACK lysing buffer, cells were cultured at 300,000 cells/cm^2^ in α-MEM medium (Dutscher) supplemented with 10% fetal bovine serum, penicillin-streptomycin, β-mercaptoethanol (50 µM, Euromedex), and osteoclastogenic cytokines including RANK-L (30 ng/mL; 310-01, PreproTech) and M-CSF (25 ng/mL).

Cells were cultured in the presence or absence of EPZ-6438 (10 µM). After approximately seven days, when multinucleated giant cells were observed, osteoclast differentiation was assessed by tartrate-resistant acid phosphatase (TRAP) staining according to the manufacturer’s protocol (387A, Sigma-Aldrich). TRAP-positive multinucleated cells were identified as osteoclasts and counted.

### Statistical analyses

Statistical analyses were performed using GraphPad Prism 8 (GraphPad Software, San Diego, CA, USA). Data distribution was assessed using the Shapiro–Wilk test. Normally distributed continuous variables are presented as mean ± SEM and were analyzed using parametric tests, whereas non-normally distributed or ordinal data are presented as median with interquartile range (IQR) and were analyzed using non-parametric tests.

For *in vitro* experiments, paired statistical tests were used whenever matched biological samples were compared between experimental conditions. For *in vivo* experiments, comparisons between independent groups were performed using unpaired statistical tests. Two-group comparisons were analyzed using paired or unpaired Student’s t-tests, or Mann–Whitney tests for non-parametric data, as appropriate. Multiple-group comparisons were performed using one-way or two-way ANOVA followed by Tukey’s multiple comparisons test, or Kruskal–Wallis tests followed by Dunn’s multiple comparisons test for non-parametric data, as appropriate. Outliers were identified using the ROUT method (Q = 1%).

For experiments using primary human cells, biological replicates corresponded to independent OA donors. For cell line experiments, biological replicates corresponded to independent experiments. Technical replicates were averaged and were not considered independent biological replicates for statistical analyses. Statistical significance was defined as p < 0.05.

## Declarations

### Ethics approval and consent to participate

Human and animal studies by the appropriate institutional review boards. The animal experiments were carried out in compliance with European directive 2010/63/EU in accordance with French legislation (decree 87/848) at the GIP CYCERON/CURB (Centre Universitaire de Ressources Biologiques, approval #D14118001). The procedures have been approved by the Ministry of Education and Research and the committee of regional ethics (CENOMEXA, France, APAFIS number #16185). Concerning human samples, the experimental protocol was approved by the local ethics committee, named “Comité de Protection des Personnes Nord Ouest III” (agreement #A13-D46-VOL.19).

### Consent for publication

Not applicable

### Availability of data and materials

The data discussed in this publication have been deposited in NCBI’s Gene Expression Omnibus (29) and are accessible through GEO Series accession number GSE300508 (https://www.ncbi.nlm.nih.gov/geo/query/acc.cgi?acc=GSE300508) or at iProX - integrated Proteome resources (Id: IPX0008212000).

### Authors’ contributions

SB, OT, CV, EM, VA, KB, CB designed research studies; SB, AV, MT, IR, BB, JP, CS, NB conducted experiments; SB, AV, MT, IR, JAL, BB, CS, NB, CP acquired data; SB, AV, MT, IR, JAL, BB, CS, OT, CV, JG, KB, CB analyzed data; BP provided reagents; SB, JAL, CB wrote the manuscript. All authors read and approved the final manuscript.

## Acknowlegments

We thank Sylvain Leclercq (Service orthopédique de la Clinique Saint-Martin, Caen France) for the femoral head samples, as well as Palma Pro and all animal facility staff (CURB, Unicaen, France) for technical assistance.

The authors declare that generative artificial intelligence tools were used as a supportive aid during manuscript preparation, including language editing, text refinement, and assistance in improving clarity and structure of the manuscript. The authors reviewed and edited the content as needed and take full responsibility for the content of the publication.

## References

1. Long H, Liu Q, Yin H, Wang K, Diao N, Zhang Y, et al. Prevalence Trends of Site-Specific Osteoarthritis From 1990 to 2019: Findings From the Global Burden of Disease Study 2019. Arthritis Rheumatol. juill 2022;74(7):1172⍰83. doi:10.1002/art.42089 PubMed PMID: 35233975; PubMed Central PMCID: PMC9543105.

2. Hunter DJ, Bierma-Zeinstra S. Osteoarthritis. The Lancet. 27 avr 2019;393(10182):1745⍰59. doi:10.1016/S0140-6736(19)30417-9 PubMed PMID: 31034380.

3. Glyn-Jones S, Palmer AJR, Agricola R, Price AJ, Vincent TL, Weinans H, et al. Osteoarthritis. The Lancet. 25 juill 2015;386(9991):376⍰87. doi:10.1016/S0140-6736(14)60802-3

4. Allas L, Boumédiene K, Baugé C. Epigenetic dynamic during endochondral ossification and articular cartilage development. Bone. 1 mars 2019;120:523⍰32. doi:10.1016/j.bone.2018.10.004

5. Caldo D, Massarini E, Rucci M, Deaglio S, Ferracini R. Epigenetics in Knee Osteoarthritis: A 2020–2023 Update Systematic Review. Life. févr 2024;14(2):2. doi:10.3390/life14020269

6. Chen L, Wu Y, Wu Y, Wang Y, Sun L, Li F. The inhibition of EZH2 ameliorates osteoarthritis development through the Wnt/β-catenin pathway. Sci Rep. 19 août 2016;6:29176. doi:10.1038/srep29176 PubMed PMID: 27539752; PubMed Central PMCID: PMC4990905.

7. Allas L, Rochoux Q, Leclercq S, Boumédiene K, Baugé C. Development of a simple osteoarthritis model useful to predict in vitro the anti-hypertrophic action of drugs. Lab Invest. janv 2020;100(1):64⍰71. doi:10.1038/s41374-019-0303-0 PubMed PMID: 31409892.

8. Allas L, Brochard S, Rochoux Q, Ribet J, Dujarrier C, Veyssiere A, et al. EZH2 inhibition reduces cartilage loss and functional impairment related to osteoarthritis. Sci Rep. 11 nov 2020;10(1):19577. doi:10.1038/s41598-020-76724-9 PubMed PMID: 33177650; PubMed Central PMCID: PMC7658239.

9. Wang J, Wang X, Ding X, Huang T, Song D, Tao H. EZH2 is associated with cartilage degeneration in osteoarthritis by promoting SDC1 expression via histone methylation of the microRNA-138 promoter. Laboratory Investigation. 1 mai 2021;101(5):600⍰11. doi:10.1038/s41374-021-00532-6

10. Miller RE, Miller RJ, Malfait AM. Osteoarthritis joint pain: the cytokine connection. Cytokine. déc 2014;70(2):185⍰93. doi:10.1016/j.cyto.2014.06.019 PubMed PMID: 25066335; PubMed Central PMCID: PMC4254338.

11. Conaghan PG, Cook AD, Hamilton JA, Tak PP. Therapeutic options for targeting inflammatory osteoarthritis pain. Nat Rev Rheumatol. juin 2019;15(6):355⍰63. doi:10.1038/s41584-019-0221-y PubMed PMID: 31068673.

12. Geraghty T, Winter DR, Miller RJ, Miller RE, Malfait AM. Neuroimmune interactions and osteoarthritis pain: focus on macrophages. Pain Rep. 2021;6(1):e892. doi:10.1097/PR9.0000000000000892 PubMed PMID: 33981927; PubMed Central PMCID: PMC8108586.

13. Neele AE, de Winther MPJ. Repressing the repressor: Ezh2 mediates macrophage activation. J Exp Med. 7 mai 2018;215(5):1269⍰71. doi:10.1084/jem.20180479 PubMed PMID: 29691302; PubMed Central PMCID: PMC5940273.

14. Adamik J, Pulugulla SH, Zhang P, Sun Q, Lontos K, Macar DA, et al. EZH2 Supports Osteoclast Differentiation and Bone Resorption Via Epigenetic and Cytoplasmic Targets. J Bone Miner Res. janv 2020;35(1):181⍰95. doi:10.1002/jbmr.3863 PubMed PMID: 31487061; PubMed Central PMCID: PMC7402427.

15. Newton MD, Swahn H, Orange DE, Lesnak JB, Price TJ, Malfait AM, et al. Cross-platform transcriptomic data integration identifies an overactive neuro-immune signature in human osteoarthritis synovium. Osteoarthritis Cartilage. avr 2026;34(4):589⍰600. doi:10.1016/j.joca.2025.08.013 PubMed PMID: 40889615; PubMed Central PMCID: PMC13112359.

16. Valenti MT, Dalle Carbonare L, Zipeto D, Mottes M. Control of the Autophagy Pathway in Osteoarthritis: Key Regulators, Therapeutic Targets and Therapeutic Strategies. International Journal of Molecular Sciences. janv 2021;22(5):5. doi:10.3390/ijms22052700

17. Wei FZ, Cao Z, Wang X, Wang H, Cai MY, Li T, et al. Epigenetic regulation of autophagy by the methyltransferase EZH2 through an MTOR-dependent pathway. Autophagy. 1 déc 2015;11(12):2309⍰22. doi:10.1080/15548627.2015.1117734 PubMed PMID: 26735435.

18. Ma Z, Wei Y, Liao T, Jie L, Yang N, Yu L, et al. Activation of vascular endothelial cells by synovial fibrosis promotes Netrin-1-induced sensory nerve sprouting and exacerbates pain sensitivity. Journal of Cellular and Molecular Medicine. 2023;27(23):3773⍰85. doi:10.1111/jcmm.17950

19. Zhang L, Li M, Li X, Liao T, Ma Z, Zhang L, et al. Characteristics of sensory innervation in synovium of rats within different knee osteoarthritis models and the correlation between synovial fibrosis and hyperalgesia. Journal of Advanced Research. 1 janv 2022;35:141⍰51. doi:10.1016/j.jare.2021.06.007

20. Rondeaux J, Groussard D, Renet S, Tardif V, Dumesnil A, Chu A, et al. Ezh2 emerges as an epigenetic checkpoint regulator during monocyte differentiation limiting cardiac dysfunction post-MI. Nat Commun. 25 juill 2023;14(1):4461. doi:10.1038/s41467-023-40186-0 PubMed PMID: 37491334; PubMed Central PMCID: PMC10368741.

21. Fang C, Qiao Y, Mun SH, Lee MJ, Murata K, Bae S, et al. Cutting Edge: EZH2 Promotes Osteoclastogenesis by Epigenetic Silencing of the Negative Regulator IRF8. J Immunol. 1 juin 2016;196(11):4452⍰6. doi:10.4049/jimmunol.1501466 PubMed PMID: 27183582; PubMed Central PMCID: PMC4875854.

22. Zhu S, Zhu J, Zhen G, Hu Y, An S, Li Y, et al. Subchondral bone osteoclasts induce sensory innervation and osteoarthritis pain. J Clin Invest. 129(3):1076⍰93. doi:10.1172/JCI121561 PubMed PMID: 30530994; PubMed Central PMCID: PMC6391093.

23. Brochard S, Boumédiene K, Mercier J, Agin V, Conrozier T, Bauge C. A single intraarticular injection of a tranexamic acid-modified hyaluronic acid (HA/TXA) alleviates pain and reduces OA development in a murine model of monosodium iodoacetate-induced osteoarthritis. Front Pharmacol. 26 août 2024;15. doi:10.3389/fphar.2024.1456495

24. Pritzker KPH, Gay S, Jimenez SA, Ostergaard K, Pelletier JP, Revell PA, et al. Osteoarthritis cartilage histopathology: grading and staging. Osteoarthritis and Cartilage. 1 janv 2006;14(1):13⍰29. doi:10.1016/j.joca.2005.07.014

25. Mankin HJ. Biochemical and metabolic aspects of osteoarthritis. Orthop Clin North Am. mars 1971;2(1):19⍰31. PubMed PMID: 4940528.

26. Brochard S, Pontin J, Bernay B, Boumediene K, Conrozier T, Baugé C. The benefit of combining curcumin, bromelain and harpagophytum to reduce inflammation in osteoarthritic synovial cells. BMC Complement Med Ther. 14 oct 2021;21(1):261. doi:10.1186/s12906-021-03435-7 PubMed PMID: 34649531; PubMed Central PMCID: PMC8515758.

27. Szklarczyk D, Kirsch R, Koutrouli M, Nastou K, Mehryary F, Hachilif R, et al. The STRING database in 2023: protein-protein association networks and functional enrichment analyses for any sequenced genome of interest. Nucleic Acids Res. 6 janv 2023;51(D1):D638⍰46. doi:10.1093/nar/gkac1000 PubMed PMID: 36370105; PubMed Central PMCID: PMC9825434.

28. Baxter EW, Graham AE, Re NA, Carr IM, Robinson JI, Mackie SL, et al. Standardized protocols for differentiation of THP-1 cells to macrophages with distinct M(IFNγ+LPS), M(IL-4) and M(IL-10) phenotypes. J Immunol Methods. mars 2020;478:112721. doi:10.1016/j.jim.2019.112721 PubMed PMID: 32033786.

29. Edgar R, Domrachev M, Lash AE. Gene Expression Omnibus: NCBI gene expression and hybridization array data repository. Nucleic Acids Res. 1 janv 2002;30(1):207⍰10. doi:10.1093/nar/30.1.207 PubMed PMID: 11752295; PubMed Central PMCID: PMC99122.

