## Supplementary material for "EZH2 coordinates inflammatory and pain-associated pathways across joint compartments in osteoarthritis": Table S1

*Table 1: List proteins deregulated by EPZ-6438 in IL-1 stimulated synoviocytes.*

| Accession | Gene name | Description | logFC | pvalue |
|---|---|---|---|---|
| P13667 | PDIA4 | Protein disulfide-isomerase A4 | -3.40093 | 0.00160207 |
| Q9P258 | RCC2 | Protein RCC2 | -3.21243 | 0.0381512 |
| P48735 | IDHP | Isocitrate dehydrogenase [NADP] mitochondrial | -0.914807 | 0.00797848 |
| Q9BZG1 | RAB34 | Ras-related protein Rab-34 | -0.775697 | 0.0230256 |
| P00568 | KAD1 | Adenylate kinase isoenzyme 1 | 0.203334 | 0.0340583 |
| P36871 | PGM1 | Phosphoglucomutase-1 | 0.212835 | 0.0101563 |
| P50395 | GDIB | Rab GDP dissociation inhibitor beta | 0.217763 | 0.0499717 |
| P30044 | PRDX5 | Peroxiredoxin-5 mitochondrial | 0.230621 | 0.0266481 |
| Q96KP1 | EXOC2 | Exocyst complex component 2 | 0.238318 | 0.0483993 |
| F5H365 | F5H365 | Protein transport protein SEC23 | 0.244969 | 0.0119121 |
| P61106 | RAB14 | Ras-related protein Rab-14 | 0.267328 | 0.0245752 |
| P18124 | RL7 | 60S ribosomal protein L7 | 0.274041 | 0.0255669 |
| O15173 | PGRC2 | Membrane-associated progesterone receptor component 2 | 0.283588 | 0.0305502 |
| P40121 | CAPG | Macrophage-capping protein | 0.287716 | 0.0112755 |
| P54578 | UBP14 | Ubiquitin carboxyl-terminal hydrolase 14 | 0.288739 | 0.0252215 |
| Q8WUM4 | PDC6I | Programmed cell death 6-interacting protein | 0.289214 | 0.0382393 |
| P49207 | RL34 | 60S ribosomal protein L34 | 0.291148 | 0.0421039 |
| P16152 | CBR1 | Carbonyl reductase [NADPH] 1 | 0.308976 | 0.0277092 |
| P62979 | RS27A | Ubiquitin-40S ribosomal protein S27a | 0.310714 | 0.00684499 |
| P20073 | ANXA7 | Annexin A7 | 0.316389 | 0.0237647 |
| A0A0C4DGS1 | A0A0C4DGS1 | Dolichyl-diphosphooligosaccharide--protein glycosyltransferase 48 kDa subunit | 0.318201 | 0.0475227 |
| P50502 | F10A1 | Hsc70-interacting protein | 0.342326 | 0.0170673 |
| Q9Y285 | SYFA | Phenylalanine--tRNA ligase alpha subunit | 0.3437 | 0.0402616 |
| Q53T59 | H1BP3 | HCLS1-binding protein 3 | 0.34453 | 0.0345412 |
| P61020 | RAB5B | Ras-related protein Rab-5B | 0.345547 | 0.0144726 |
| P18084 | ITB5 | Integrin beta-5 | 0.349897 | 0.0223674 |
| Q5JR08 | Q5JR08 | Rho-related GTP-binding protein RhoC (Fragment) | 0.350433 | 0.0467418 |
| P52597 | HNRPF | Heterogeneous nuclear ribonucleoprotein F | 0.351871 | 0.00952878 |
| P26640 | SYVC | Valine--tRNA ligase | 0.360656 | 0.0143028 |
| O15118 | NPC1 | NPC intracellular cholesterol transporter 1 | 0.368992 | 0.020931 |
| E9PLL6 | E9PLL6 | 60S ribosomal protein L27a | 0.369018 | 0.0198409 |
| Q9H3U1 | UN45A | Protein unc-45 homolog A | 0.374484 | 0.0325441 |
| P01023 | A2MG | Alpha-2-macroglobulin | 0.388073 | 0.0199562 |
| P58335 | ANTR2 | Anthrax toxin receptor 2 | 0.402749 | 0.0458711 |
| B8ZZQ6 | B8ZZQ6 | Prothymosin alpha | 0.40537 | 0.0257504 |
| A2A274 | A2A274 | Aconitate hydratase mitochondrial | 0.419552 | 0.0157098 |
| P32969 | RL9 | 60S ribosomal protein L9 | 0.422249 | 0.00070824 |
| P62826 | RAN | GTP-binding nuclear protein Ran | 0.429566 | 0.0358472 |
| P07108 | ACBP | Acyl-CoA-binding protein | 0.429875 | 0.00098358 |
| P52895 | AK1C2 | Aldo-keto reductase family 1 member C2 | 0.437757 | 0.0206759 |
| O00499 | BIN1 | Myc box-dependent-interacting protein 1 | 0.438221 | 0.0301449 |
| P30050 | RL12 | 60S ribosomal protein L12 | 0.446763 | 0.0275765 |
| O94832 | MYO1D | Unconventional myosin-Id | 0.454851 | 0.0292924 |
| P04080 | CYTB | Cystatin-B | 0.462475 | 0.0247684 |
| Q13442 | HAP28 | 28 kDa heat- and acid-stable phosphoprotein | 0.469984 | 0.0142197 |
| P22695 | QCR2 | Cytochrome b-c1 complex subunit 2 mitochondrial | 0.474312 | 0.0353184 |
| Q14195 | DPYL3 | Dihydropyrimidinase-related protein 3 | 0.480645 | 0.0335528 |
| Q16401 | PSMD5 | 26S proteasome non-ATPase regulatory subunit 5 | 0.487153 | 0.0105066 |
| O95202 | LETM1 | Mitochondrial proton/calcium exchanger protein | 0.48747 | 0.033909 |
| Q9UHV9 | PFD2 | Prefoldin subunit 2 | 0.491241 | 0.0287517 |
| A0A0A6YYA0 | A0A0A6YYA0 | Protein TMED7-TICAM2 | 0.491629 | 0.0328677 |
| E7EPK1 | E7EPK1 | Septin | 0.494426 | 0.038302 |
| A0A1W2PQT3 | A0A1W2PQT3 | Malic enzyme | 0.498047 | 0.00788401 |
| P32321 | DCTD | Deoxycytidylate deaminase | 0.502155 | 0.0411568 |
| P61923 | COPZ1 | Coatomer subunit zeta-1 | 0.532699 | 0.0439205 |
| A0A0A0MS51 | A0A0A0MS51 | Actin-depolymerizing factor | 0.533591 | 0.0391932 |
| Q00839 | HNRPU | Heterogeneous nuclear ribonucleoprotein U | 0.533669 | 0.031206 |
| P09497 | CLCB | Clathrin light chain B | 0.536462 | 0.0376439 |
| P08253 | MMP2 | 72 kDa type IV collagenase | 0.539859 | 0.0159 |
| P07204 | TRBM | Thrombomodulin | 0.544198 | 0.0312993 |
| Q15046 | SYK | Lysine--tRNA ligase | 0.544365 | 0.0284348 |
| A0A1C7CYX9 | A0A1C7CYX9 | Dihydropyrimidinase-related protein 2 | 0.545378 | 0.00983617 |
| P17931 | LEG3 | Galectin-3 | 0.548639 | 0.0268366 |
| Q15717 | ELAV1 | ELAV-like protein 1 | 0.553402 | 0.00215299 |

| | | | | |
|---|---|---|---|---|
| Q99536 | VAT1 | Synaptic vesicle membrane protein VAT-1 homolog | 0.561105 | 0.0390699 |
| P11766 | ADHX | Alcohol dehydrogenase class-3 | 0.56493 | 0.0103023 |
| Q9NX46 | ADPRS | ADP-ribose glycohydrolase ARH3 | 0.573112 | 0.0420053 |
| P55036 | PSMD4 | 26S proteasome non-ATPase regulatory subunit 4 | 0.59972 | 0.0498809 |
| P27105 | STOM | Stomatin | 0.618933 | 0.0477259 |
| O95295 | SNAPN | SNARE-associated protein Snapin | 0.624175 | 0.0355106 |
| P07900 | HS90A | Heat shock protein HSP 90-alpha | 0.63659 | 0.00174499 |
| Q9NZU5 | LMCD1 | LIM and cysteine-rich domains protein 1 | 0.636793 | 0.0483049 |
| P20742 | PZP | Pregnancy zone protein | 0.654117 | 0.0137395 |
| Q14444 | CAPR1 | Caprin-1 | 0.686128 | 0.0423988 |
| P60866 | RS20 | 40S ribosomal protein S20 | 0.68851 | 0.0207338 |
| J3QRI7 | J3QRI7 | 60S ribosomal protein L26 (Fragment) | 0.698098 | 0.0417818 |
| P42677 | RS27 | 40S ribosomal protein S27 | 0.72299 | 0.0350072 |
| P12268 | IMDH2 | Inosine-5'-monophosphate dehydrogenase 2 | 0.741286 | 0.0447653 |
| P62829 | RL23 | 60S ribosomal protein L23 | 0.742774 | 0.0441349 |
| A0A7P0Z4A2 | A0A7P0Z4A2 | Sorting nexin-9 | 0.755497 | 0.0424821 |
| Q8N5K1 | CISD2 | CDGSH iron-sulfur domain-containing protein 2 | 0.767913 | 0.00676764 |
| Q13263 | TIF1B | Transcription intermediary factor 1-beta | 0.769998 | 0.0401198 |
| P48426 | PI42A | Phosphatidylinositol 5-phosphate 4-kinase type-2 alpha | 0.770961 | 0.0340927 |
| A0A087WY55 | A0A087WY55 | Chromosome 6 open reading frame 55 isoform CRA_b | 0.772326 | 0.0324466 |
| Q8WWM9 | CYGB | Cytoglobin | 0.774351 | 0.0109495 |
| A0A2R8Y484 | A0A2R8Y484 | Integrin-associated protein (Fragment) | 0.774624 | 0.0282902 |
| Q9Y2A7 | NCKP1 | Nck-associated protein 1 | 0.779765 | 0.0488668 |
| P41226 | UBA7 | Ubiquitin-like modifier-activating enzyme 7 | 0.789098 | 0.0454478 |
| E9PNW4 | E9PNW4 | CD59 glycoprotein | 0.789828 | 0.0113317 |
| E5RGS4 | E5RGS4 | Prefoldin subunit 1 | 0.79671 | 0.0163729 |
| Q9NT62 | ATG3 | Ubiquitin-like-conjugating enzyme ATG3 | 0.800326 | 0.0452554 |
| Q9BVC6 | TM109 | Transmembrane protein 109 | 0.805061 | 0.00516605 |
| E9PH64 | E9PH64 | NADH dehydrogenase [ubiquinone] 1 beta subcomplex subunit 9 | 0.807782 | 0.0439846 |
| P23381 | SYWC | Tryptophan--tRNA ligase cytoplasmic | 0.812357 | 0.0415376 |
| A0A2R8Y7R2 | A0A2R8Y7R2 | Hemoglobin subunit beta | 0.861646 | 0.00354612 |
| P36405 | ARL3 | ADP-ribosylation factor-like protein 3 | 0.890901 | 0.0289507 |
| Q9Y265 | RUVB1 | RuvB-like 1 | 0.895786 | 0.0269516 |
| P46976 | GLYG | Glycogenin-1 | 0.907195 | 0.0316347 |
| P55735 | SEC13 | Protein SEC13 homolog | 0.935264 | 0.0292658 |
| P80723 | BASP1 | Brain acid soluble protein 1 | 0.95867 | 0.0193001 |
| P61201 | CSN2 | COP9 signalosome complex subunit 2 | 0.984665 | 0.0150697 |
| Q2TAA2 | IAH1 | Isoamyl acetate-hydrolyzing esterase 1 homolog | 1.0192 | 0.00271183 |
| Q8NDC0 | MISSL | MAPK-interacting and spindle-stabilizing protein-like | 1.02822 | 0.0419808 |
| Q96AT9 | RPE | Ribulose-phosphate 3-epimerase | 1.03937 | 0.0471483 |
| D6RFH4 | D6RFH4 | Cytochrome b5 type B | 1.07564 | 0.0476349 |
| P50281 | MMP14 | Matrix metalloproteinase-14 | 1.10562 | 0.0462285 |
| Q5T7C4 | Q5T7C4 | High mobility group protein B1 | 1.10696 | 0.044598 |
| P02795 | MT2 | Metallothionein-2 | 1.1112 | 0.0253598 |
| Q9BPX5 | ARP5L | Actin-related protein 2/3 complex subunit 5-like protein | 1.16067 | 0.00224028 |
| Q9GZT8 | NIF3L | NIF3-like protein 1 | 1.16915 | 0.00787084 |
| A0A0C4DGV4 | A0A0C4DGV4 | Late endosomal/lysosomal adaptor and MAPK and MTOR activator 5 | 1.18292 | 0.0463546 |
| A0A3B3IRS7 | A0A3B3IRS7 | Lysosomal cobalamin transport escort protein LMBD1 | 1.22317 | 0.0130033 |
| Q5H9A7 | Q5H9A7 | Metalloproteinase inhibitor 1 | 1.22839 | 0.0209772 |
| P21817 | RYR1 | Ryanodine receptor 1 | 1.2339 | 0.0305715 |
| P48643 | TCPE | T-complex protein 1 subunit epsilon | 1.26143 | 0.0342727 |
| Q6IBS0 | TWF2 | Twinfilin-2 | 1.28027 | 0.0435127 |
| P09619 | PGFRB | Platelet-derived growth factor receptor beta | 1.3079 | 0.0286426 |
| P51151 | RAB9A | Ras-related protein Rab-9A | 1.32109 | 0.0429648 |
| P02545 | LMNA | Prelamin-A/C | 1.44821 | 0.0455913 |
| Q00653 | NFKB2 | Nuclear factor NF-kappa-B p100 subunit | 1.45501 | 0.00848168 |
| P17813 | EGLN | Endoglin | 1.52678 | 0.0353231 |
| O60547 | GMDS | GDP-mannose 4 6 dehydratase | 1.54079 | 0.0320901 |
| Q14165 | MLEC | Malectin | 1.5523 | 0.0359763 |
| A0A7I2V5M3 | A0A7I2V5M3 | Procathepsin L | 1.68857 | 0.0410056 |
| Q07955 | SRSF1 | Serine/arginine-rich splicing factor 1 | 1.84574 | 0.0347603 |
| P69905 | HBA | Hemoglobin subunit alpha | 2.06381 | 0.0288423 |
| O94874 | UFL1 | E3 UFM1-protein ligase 1 | 2.29529 | 0.00484066 |
| P62070 | RRAS2 | Ras-related protein R-Ras2 | 2.34482 | 0.0169834 |
