## Supplementary material for "EZH2 coordinates inflammatory and pain-associated pathways across joint compartments in osteoarthritis": figures S1, S2, S3

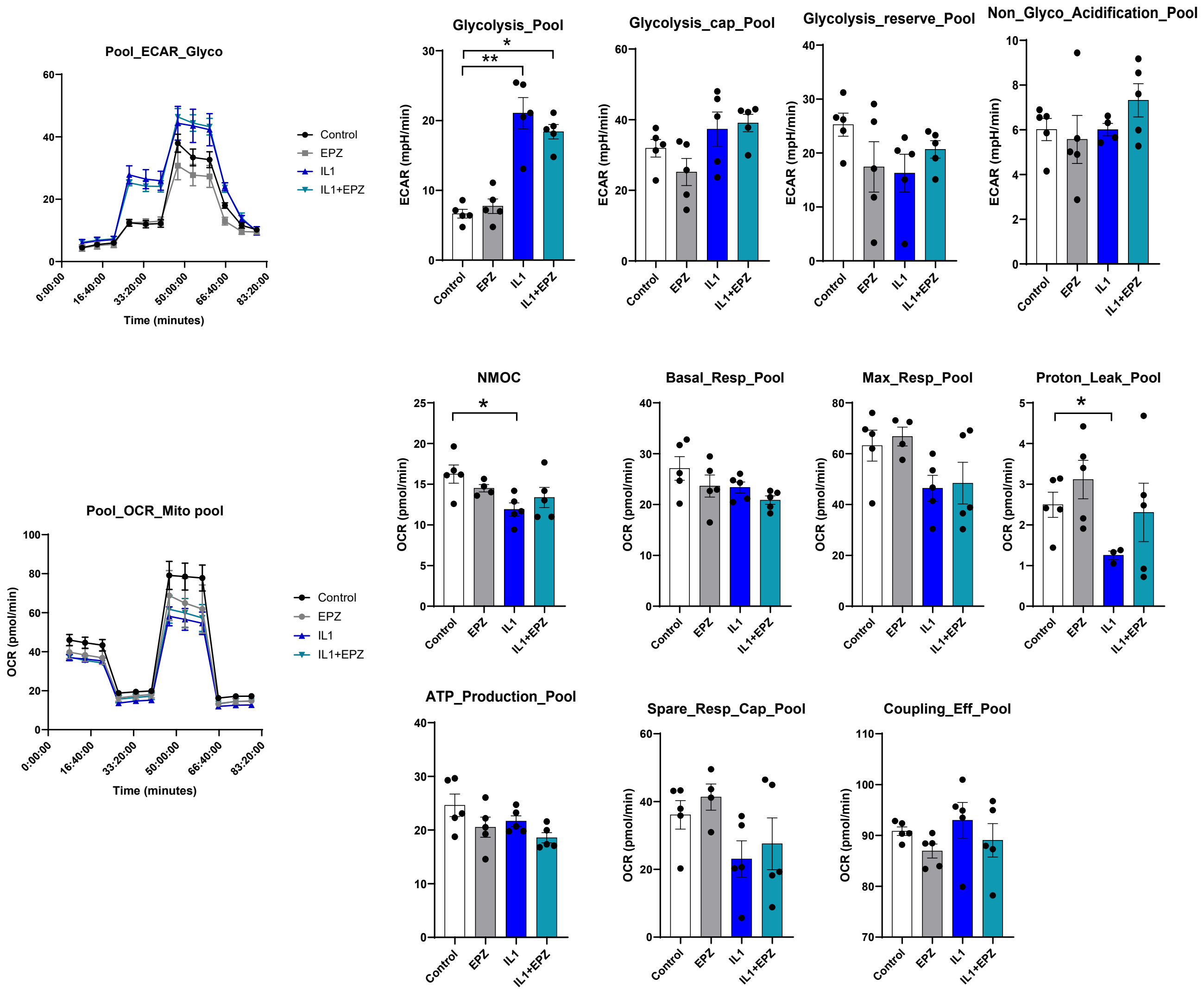


**Figure S1: Seahorse analysis of metabolic parameters following EZH2 inhibition in OA fibroblast-like synoviocytes.**

Fibroblast-like synoviocytes were stimulated with IL-1β (1 ng/ml) in the presence or absence of EPZ-6438 (10 μM). After 24 h of treatment, glycolytic and mitochondrial activities were assessed by Seahorse extracellular flux analysis. Extracellular acidification rate (ECAR) and oxygen consumption rate (OCR) were measured every 5 minutes before and after sequential addition of metabolic modulators. Data were analyzed using Wave software. Experiments were independently repeated at least three times. Data are presented as mean ± SEM. Statistical significance was defined as *p < 0.05 and **p < 0.01.

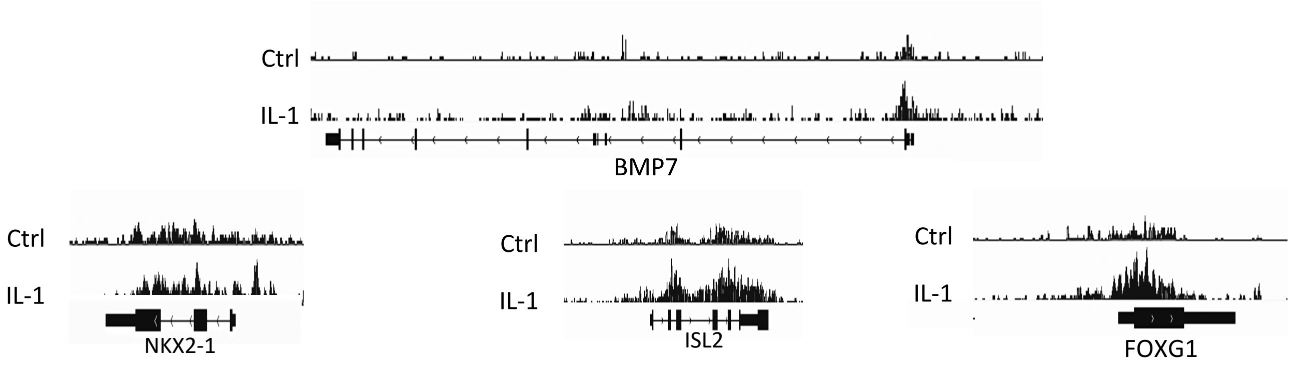


**Figure S2: EZH2 occupancy at promoters of genes associated with neurogenesis and axon guidance pathways.** Fibroblast-like synoviocytes were stimulated with IL-1β for 48 h, and EZH2 chromatin occupancy was assessed by ChIP-seq analysis (as described in Figure 5). A binary comparison of ChIP-seq peaks was performed to identify common and condition-specific EZH2-bound regions. Representative EZH2-enriched peaks located at promoter regions of BMP7, NKX2-1, ISL2, and FOXG1, genes associated with neurogenesis and axon guidance processes, are shown.

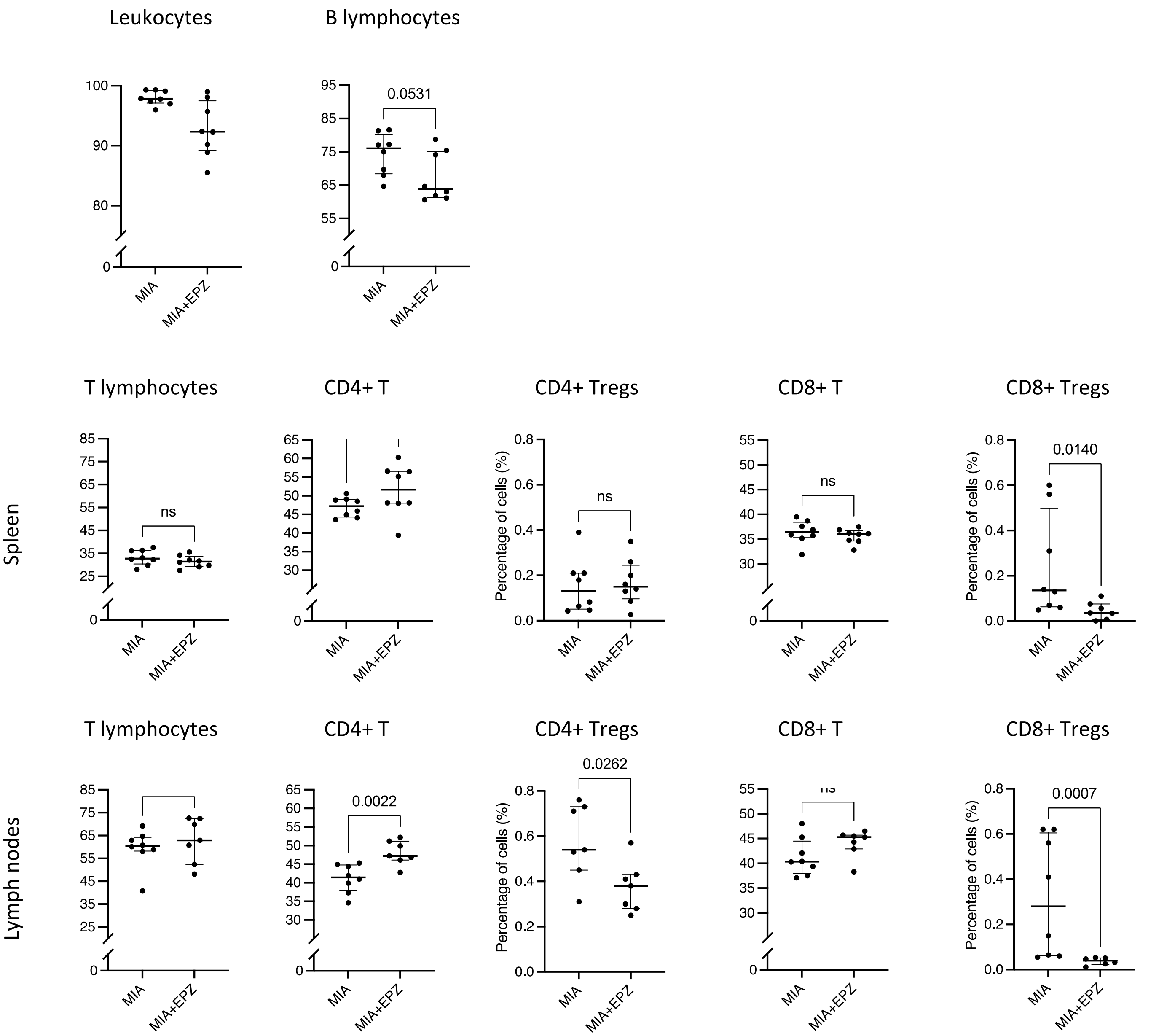


**Figure S3: Flow cytometric characterization of lymphocyte populations in spleen and lymph nodes of OA mice following EZH2 inhibition.**

Leukocyte and lymphocyte subsets were analyzed by flow cytometry in spleen and lymph nodes from OA mice treated with vehicle or EPZ-6438. (a) Percentage of total leukocytes and B lymphocytes in the spleen. (b) Frequency of total T lymphocytes, $CD4^+$ T cells, $CD4^+$ regulatory T cells (Tregs), $CD8^+$ T cells, and $CD8^+$ Tregs in the spleen. (c) Frequency of total T lymphocytes, $CD4^+$ T cells, $CD4^+$ Tregs, $CD8^+$ T cells, and $CD8^+$ Tregs in lymph nodes. Cells were isolated and stained with appropriate surface markers for flow cytometry analysis. Samples were acquired using a FACSVerse cytometer and analyzed with FlowJo software. Results are presented as median ± interquartile range (n = 8 mice). Outliers were identified using the ROUT method (Q = 1%) and may have been excluded, potentially resulting in different numbers of analyzed samples. Each dot represents an individual mouse. Statistical significance was assessed using the Mann–Whitney test; $p < 0.05$ was considered statistically significant.
